# A diversified, widespread microbial gene cluster encodes homologs of methyltransferases involved in methanogenesis

**DOI:** 10.1101/2023.07.31.551370

**Authors:** Duncan J. Kountz, Emily P. Balskus

**Affiliations:** Department of Chemistry and Chemical Biology, Harvard University, Cambridge, Massachusetts 02138, United States; Howard Hughes Medical Institute, Harvard University, Cambridge, Massachusetts 02138, United States

**Author notes:** Author information: Duncan J. Kountz; Emily P. Balskus.

**Keywords:** Corrinoid methyltransferases, methanogenesis, selenium metabolism, sulfur metabolism, bacteria

## Abstract

Analyses of microbial genomes have revealed unexpectedly wide distributions of enzymes from specialized metabolism, including methanogenesis, providing exciting opportunities for discovery. Here, we identify a family of gene clusters (the type 1 *mlp* gene clusters (MGCs)) that encodes homologs of the soluble coenzyme M methyltransferases (SCMTs) involved in methylotrophic methanogenesis and is widespread in bacteria and archaea. Type 1 MGCs are expressed and regulated in medically, environmentally, and industrially important organisms, making them likely to be physiologically relevant. Enzyme annotation, analysis of genomic context, and biochemical experiments suggests these gene clusters play a role in methyl-sulfur and/or methyl-selenide metabolism in numerous anoxic environments, including the human gut microbiome, potentially impacting sulfur and selenium cycling in diverse, anoxic environments.

## Introduction

Other the past 50 years, biochemists and microbiologists have made much progress in elucidating the enzymology of methanogenesis (1,2). This reservoir of biochemical knowledge presents the opportunity to identify homologs of methanogen specific enzymes in organisms that are categorically not methanogens. In doing so, we may discover previously unappreciated metabolic pathways with global or ecological relevance. Moreover, studying homologs of methanogen-specific enzymes in non-methanogenic organisms could shed light on the evolution of methanogenic pathways and organisms as well as the potential adaptation of methanogen genes to perform novel functions.

Methanogenesis can be regarded as a specialized metabolism—its pathways are biochemically distinct from those of most organisms and not expected to interface with other metabolic processes when horizontally transferred. However, methanogen cofactors and homologs of methanogen enzymes are indeed found in non-methanogens. Examples include the use of methanofuran- and methanopterin-dependent enzymes in aerobic methylotrophy (3), deazaflavin redox cofactor F_420_ in bacterial metabolism (4), coenzyme M in alkene catabolism (5), heterodisulfide reductase-like enzymes in various metabolisms, including sulfate reduction (6), acetogenesis (7), and the degradation of benzoate (8). In each case the “methanogen” enzymes are performing an interesting, novel, and important function.

We wondered if this logic extends to the soluble coenzyme M methyltransferases (SCMTs) from methylotrophic methanogenesis. SCMTs are corrinoid-dependent methyltransferases that transfer a methyl group from a methylated corrinoid-binding protein (corrinoid protein) to coenzyme M to form methyl-coenzyme M, the immediate precursor of methane (Fig. 1) (9). SCMTs belong to the very large and sequence-diverse uroporphyrinogen decarboxylase (UroD) superfamily, which includes uroporphyrinogen decarboxylase, cobalamin-independent methionine synthase, epoxide coenzyme M transferase, and a variety of corrinoid methyltransferases including *O*-, *S*-, and halogen-demethylases (10–16). However, we focused on proteins having >25% amino acid identity to a characterized SCMT. We call these enzymes <u>M</u>taA-like <u>p</u>roteins, or Mlps, after MtaA, the methanol-specific SCMT in *Methanosarcina* species (16). To our knowledge, these Mlps have not been assigned a role outside of methanogens.

**Fig. 1.**
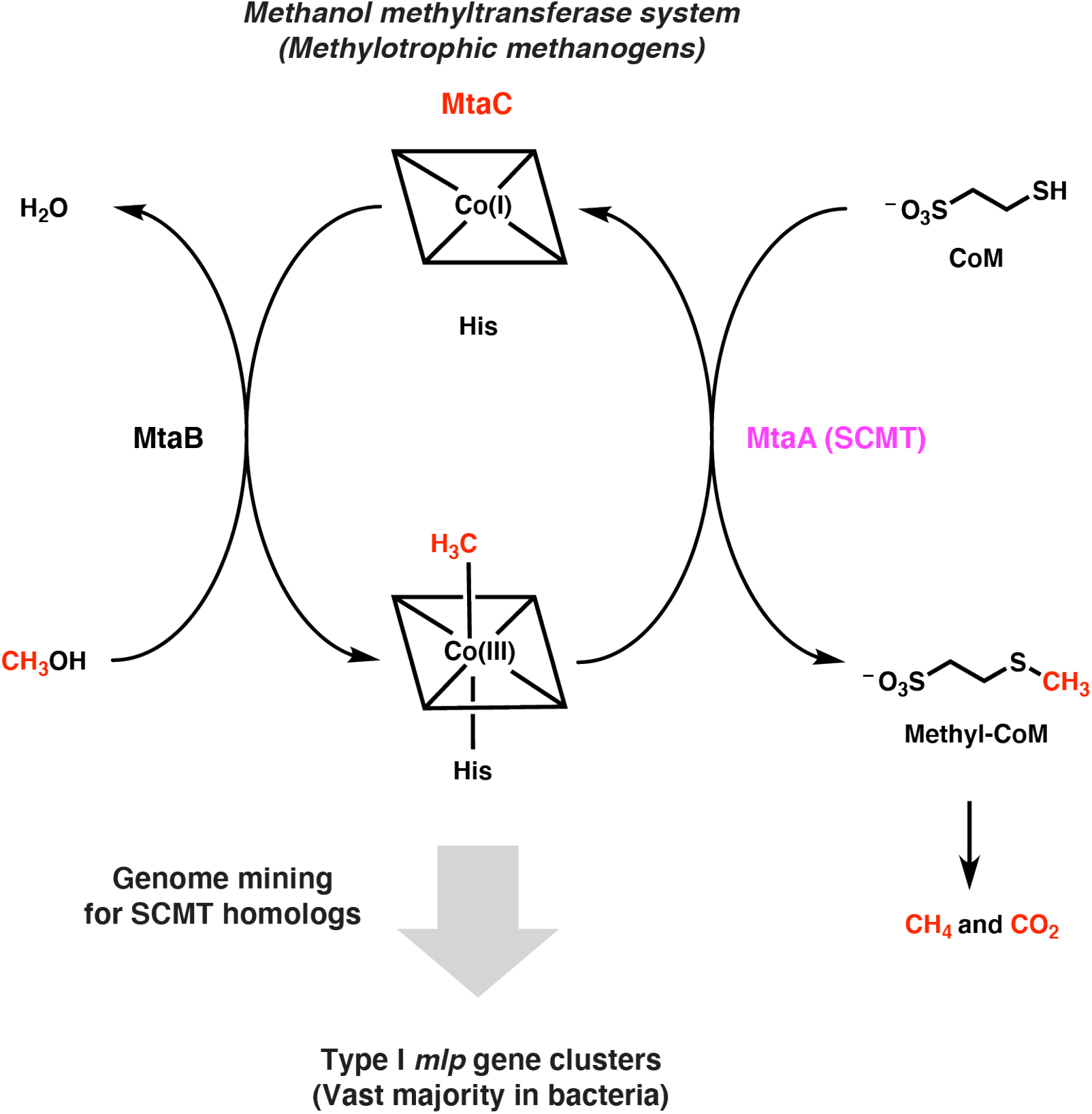
Searching genomes for the soluble coenzyme M methyltransferase (SCMT) MtaA involved in methylotrophic methanogenesis identifies related enzymes in non-methanogens. Overview of genome mining strategy used in this work. The methanol-specific methanol:corrinoid methyltransferase MtaB transfers the methyl group of methanol to the cognate corrinoid protein MtaC (the corrinoid cofactor is represented by a square containing a cobalt ion). The SCMT MtaA then transfers to the methyl group from MtaC to coenzyme M (CoM). The methyl group incorporated onto CoM is then disproportionated into methane and carbon dioxide or solely reduced to methane, depending on the organism and environmental conditions.

Here, we propose a prospective role for Mlps in a wide range of microorganisms by identifying a diverse, widespread, and homologous set of gene clusters that encode multiple Mlps, a corrinoid protein and a corrinoid reductive activase (Ram/RACE enzyme). Gene cluster annotation, bioinformatic analysis, and biochemical logic suggests that the encoded enzymes have two functions—methyl-selenide recycling and/or methionine salvage—depending on the organism. These results illustrate the utility of using known enzymes involved in metabolisms of interest to discover uncharacterized microbial gene clusters that likely have intriguing and ecologically important functions.

## Materials and methods

### General methods and procedures

Manipulations and visualizations of genomic data were performed using Geneious Prime 2020.0.4 software (17). BLASTP searches were conducted through the Geneious interface using either National Center for Biotechnology Information (NCBI) nr/rt database or custom databases generated by the user. In all cases we used the BLOSUM62 matrix and a gap opening penalty of 11 and a gap extension penalty of 1. Protein alignments we performed using MUSCLE (18).

### Identification of *mlp* gene clusters

To identify homologs of SCMTs in *C. ljungdahlii*, we generated a protein database of all the predicted proteins encoded in the *C. ljungdahlii* DSM 13528 genome, and performed at BLASTP search of this database using the *Methanosarcina barkeri* MtbA protein sequence (Swiss-Prot ID: O30640) as a query. We then inspected the genomic context of the all the hits with E-values less than 1.0e-10.

To identify additional MGCs, we performed BLASTP searches against NCBI’s nr/nt database using *C. ljungdahlii* Mlp1, Mlp2, and Mlp3 (locus tags in Table S1) as queries. We collected the top 1000 hits from each BLAST search. For the Mlp1 search, this yielded hits with as low as 19.8% amino acid identity to the query. For the Mlp2 and Mlp3 searches, the minimum percent identities of the hits were 28.0% and 24.9%, respectively. We then manually inspected the genomic context of each hit to determine whether they were found in true MGCs. A “true” MGC met the following criteria: 1) it encodes at least two Mlps, a corrinoid protein, and a Ram protein, 2) each of these components are transcribed in the same direction with no intervening divergently-transcribed genes, 3) there is no more than 2 kb of DNA in between the genes noted in 1, and 4) the gene cluster encodes no homologs of tetrahydrofolate methylases within 20 kb of any of the genes noted in 1. Using these methods and definitions, we identified 367 MGCs.

### SSN analysis of MGC proteins

We generated all SSNs using the Enzyme Function Initiative Enzyme Similarity Tool (EFI-EST) webtool (19). For each SSN, we first compiled the sequences of characterized superfamily members and superfamily members that are encoded in the 367 MGCs identified above, generating a superfamily-specific “preliminary input list”. We then pruned each list to ensure that only quality sequences (those lacking frameshifts or internal stops) within a certain length window were included. For the Mlp SSN, length window was 200 to 470 amino acids. The resulting sequence list was termed the “input list”. We next input both the appropriate superfamily and input list into the EFI-EST web server. For the Mlp SSN, the superfamily was IPR027980, for the Ram/RACE SSN, it was IPR027980, and for the corrinoid protein SSN it was IPR006158. We selected alignment score thresholds such that most MGC proteins fell into common network clusters, but were still as separated as possible from non-MGC proteins. We selected node representation percent identities to the highest that consistent with efficient use of the Cytoscape GUI on an ordinary laptop MacIntosh computer. For each SSN, we used the UniRef90 option and otherwise default parameters. Once generated, the SSNs were visualized with Cytoscape software (version 3.8.0) using the “preferred” layout (20).

### Phylogenetic analysis of *mlp* gene cluster proteins

We compiled proteins encoded by type I MGCs and sequences of related characterized proteins and then aligned the resulting protein sequences using MUSCLE with default parameters, and trimmed the alignments manually. All trees were generated using FastTree (approximately-maximum likelihood method) with default parameters (21).

We generated the rooted phylogenetic tree of the Mlps as follows. We first compiled the sequences of Mlps1, Mlps2, and Mlps3 from type I MGCs included in the above SSN. We added the SCMTs identified in the unrooted tree to this list, as well as the known phenolic *O*-demethylases OdmB and VdmB (UniProt IDs: O87604 and Q6W001, respectively). The SCMTs were identified as the smallest set of branches containing the characterized SCMTs MtaA, MtbA, and MtsA (all from *Methanosarcina barkeri* Fusaro), but excluding the Mlps3. The SCMTs were identified as the smallest clade in the unrooted tree which contained the known SCMTs MtaA, MtbA, and MtsA. The tree was rooted on the branch connecting the *O*-demethylases to the rest of the sequences, based on the hypothesis that they serve as an appropriate outgroup.

## Results

While manually inspecting the genome of *Clostridium ljungdahlii* DSM 13528, a potential producer of biofuels, we noticed that this organism encodes multiple members of the UroD superfamily. To see if these genes are closely related to SCMTs, we performed BLASTP searches of the SCMT MtbA (*Methanosarcina barkeri* Fusaro, *Mb*MtbA) against the predicted *C. lungdahlii* DSM 13528 proteome. We identified seven hits with an E value < 1e-09 (Table S1). The highest hit was CLJU_RS09375, which showed 32.5% amino acid (aa) identity to *Mb*MtbA. It is worth noting that the SCMTs MtbA, MtaA, and MtsA (all from *M. barkeri* Fusaro) show only about 30% aa identity to each other, suggesting that CLJU_RS09375 could be a close relative of the SCMTs. CLJU_RS09375 had only 17.6% and 17.0% aa identity to the UroD superfamily phenolic *O*-demethylases OdmB and VdmB, respectively.

To explore the physiological role of CLJU_RS09375, we next examined its genomic context. CLJU_RS09375 is encoded in a gene cluster that includes genes encoding a corrinoid protein (CLJU_RS09370), a “bacterial-type” corrinoid protein reductive activase (Ram/RACE protein) (CLJU_RS09380), and two additional Mlps (CLJU_RS09360 and CLJU_RS09365) (Fig. 2a). This gene cluster has been independently predicted to form an operon by OperonDB, an automatic operon prediction database (26). Because this gene cluster encodes multiple Mlps, we named it the *mlp* gene cluster (MGC). We named the three Mlps in the gene cluster Mlp1, Mlp2, and Mlp3, (specifically, *Cl*Mlp1, 2, and 3) in order of their appearance in the gene cluster from the presumed transcriptional start site (see Fig. 2a).

**Fig. 2.**
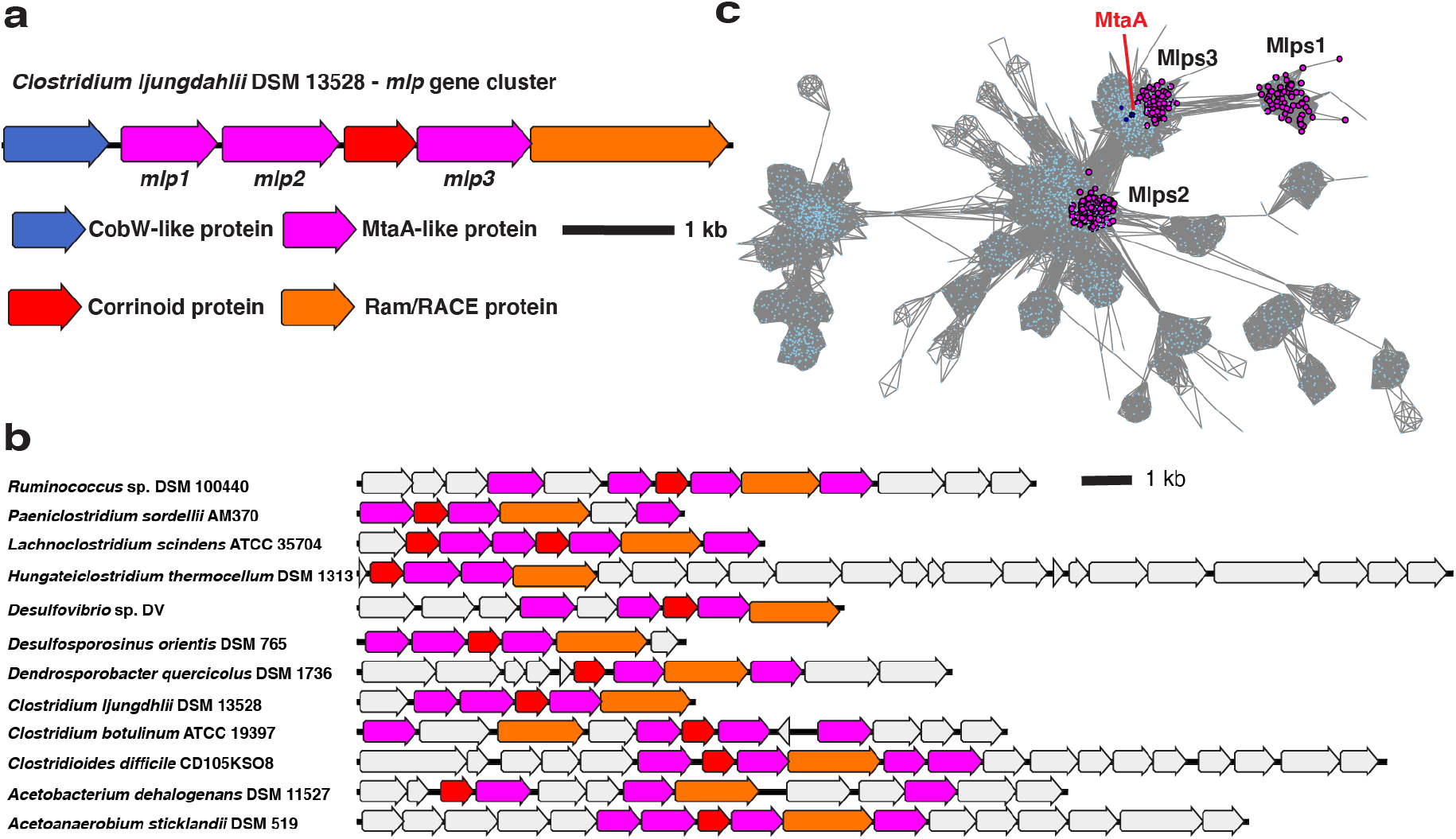
Mlp gene clusters (MGCs) are similar in genetic composition and protein seqence. **a** The MGC from *C. ljungdahlii*. **b** Representative type I MGC. Coloring of the core MGC genes is the same as in a, but non-core genes are colored grey. **c** A protein SSN of the UroD superfamily (IPR000257) showing the clustering of Mlps1, 2, and 3. Nodes representing MtsA, MtbA, and MtaA (all from *M. barkeri* Fusaro) are shown in dark blue. The alignment score cutoff was 40 and the node representation percent cut was 50 % aa identity. The SSN shown is a trimmed version of the full SSN, with all nodes removed that are neither directly or indirectly connected to the Mlps1, 2, or 3.

Given the relatively high sequence similarity of the MGC Mlps to SCMTs, it seems reasonable to hypothesize that these enzymes are corrinoid methyltransferases. This is consistent with the presence of genes encoding a corrinoid protein and a Ram/RACE protein in the MGC. The former is a methyl group carrier (27), while the latter is an ATP-dependent corrinoid protein reductase that reductively activates the corrinoid protein to its catalytically-active Co(I) state (28,29). Corrinoid methyltransferases, corrinoid proteins, and Ram/RACE proteins are frequently colocalized (30–32). In bacteria, such gene clusters are involved in methylotrophic metabolism and participate in transferring a methyl group from a donor substrate to tetrahydrofolate (H_4_folate) via a methyl-corrinoid protein intermediate, a reaction catalyzed by a methyl-corrinoid protein:H_4_folate methyltransferase (15,30). In most of the bacterial methyltransferase pathways examined to date, the H_4_folate methylase is encoded in the same gene cluster as the corrinoid protein and methyl donor methyltransferase (30,31). It is therefore surprising that H_4_folate methyltransferase is missing from the *C. ljungdahlii* MGC. While it is possible that a H_4_folate methyltransferase encoded elsewhere in the *C. ljungdahlii* genome could cooperate with the MGC methyltransferase system, it is also possible that H_4_folate is not involved or that one of the MGC Mlps is an undiscovered H_4_folate methyltransferase.

In parallel, we located similar gene clusters in *C. sporogenes* DSM 795 and *Hungateiclostridium* (*Clostridium*) *thermocellum* DSM 1313. Like the MGC from *C. ljungdahlii*, the *C. sporogenes* and *H. thermocellum* gene clusters encode multiple Mlps (each with about 30% aa identity to the SCMTs), a corrinoid protein, and a Ram/RACE protein (Fig. 2b). These gene clusters also lack a H_4_folate methylase. However, unlike *C. ljungdahlii*, which encodes H_4_folate methylases elsewhere in its genome, homologs of H_4_folate methylases are absent from the *C. sporogenes* and *H. thermocellum* genomes, suggesting that H_4_folate is not involved in the pathways encoded by these gene clusters. Because of the similarity of the *C. sporogenes* and *H. thermocellum* gene clusters to the *C. ljungdahlii* MGC, we refer to then as the *Cs*MGC, *Ht*MGC, and *Cl*MGC, respectively.

We next wondered if related MGCs are found in additional organisms. To address this, we BLASTed *Cl*Mlp1, 2, and 3 against the NCBI non-redundant protein database. Many of the organisms that had high alignment score hits for *Cl*Mlp1 also encoded high alignment score hits for *Cl*Mlp2 and 3. When we examined the genomic contexts of these hits, we identified gene clusters that resemble the *C. ljungdahlii* MGC (Fig. 2b).

We next asked if these MGCs are homologous or analogous. Because components of a corrinoid methyltransferase pathway are commonly encoded in gene clusters (14,15,30–32), evolutionary pressure to colocalize corrinoid methyltransferase genes could have resulted in superficially similar MGCs that share no evolutionary relationship. Alternatively, these gene clusters could be descended from a common ancestor (i.e. are homologous), which would also suggested a shared function in diverse organisms. To address this question, we identified 367 organisms encoding likely MGCs (see materials and methods). We next constructed several protein sequence similarity networks (SSNs) (19), reasoning that if MGCs are homologous, each conserved component should form its own cluster in an SSN of the protein family to which that component belongs. Because there are multiple *mlp* genes in each MGC, there are two SSN analysis outcomes that are consistent with the homology hypothesis. First, all Mlps from the MGCs could be grouped within a single cluster in the SSN, which would be expected if the multiple *mlp* genes in each MGC arose from gene duplication. Second, the MGC Mlps could each represent different lineages and would therefore likely form distinct clusters within an SSN.

We first constructed an SSN (Fig. 2c) of the IPR000257 superfamily, which includes the Mlps. We found that 80.9% of the MGC Mlps localized into one of three sub-clusters within a larger cluster (Fig. 2c). Interestingly, these three sub-clusters correspond to the three Mlps from the *C. ljungdahlii* MGC, with *Cl*Mlp1, 2, and 3 being grouped into distinct sub-clusters. Inspection of other MGCs reveals that almost all encode Mlps from clusters 2 and 3, while a gene encoding an Mlp from cluster 1 was present a majority of the time, but is often absent. We denote the proteins in the sub-cluster containing *Cl*Mlp1 as Mlps1, and those from sub-clusters containing *Cl*Mlp2 and 3 as Mlps2, and Mlps3, respectively. Altogether, this analysis shows that a large number of MGCs encode at least an Mlp2 and an Mlp3. We designate such gene clusters as “type 1 MGCs”.

Because >95% of MGCs identified appear to be type 1 MGCs, we selected these MGCs for further study. The fact that the type 1 MGC Mlps fall into three separate clusters in the SSN suggests they might represent three distinct Mlp lineages, rather than a single lineage that arose from recent gene duplication events. To solidify this inference, we constructed a phylogenetic tree of the Mlps and other members of the UroD superfamily that are close relatives of known corrinoid methyltransferases (Figure 3a). The Mlps 1, 2, and 3 are situated in a large branch that includes the SCMTs, to the exclusion of other known UroD superfamily corrinoid methyltransferases such as the aromatic *O*-demethylases (13). When rooted (Fig. 3b), we found that the Mlps 1, 2 and 3 formed distinct, well-supported clades. Interestingly, the Mlps2 form a sister clade to the clade containing the SCMTs and the Mlps1 and Mlps3. Within this clade, the Mlps1 and Mlps3 form sister clades emerging from the SCMTs, which therefore appear to be paraphyletic.

**Fig. 3.**
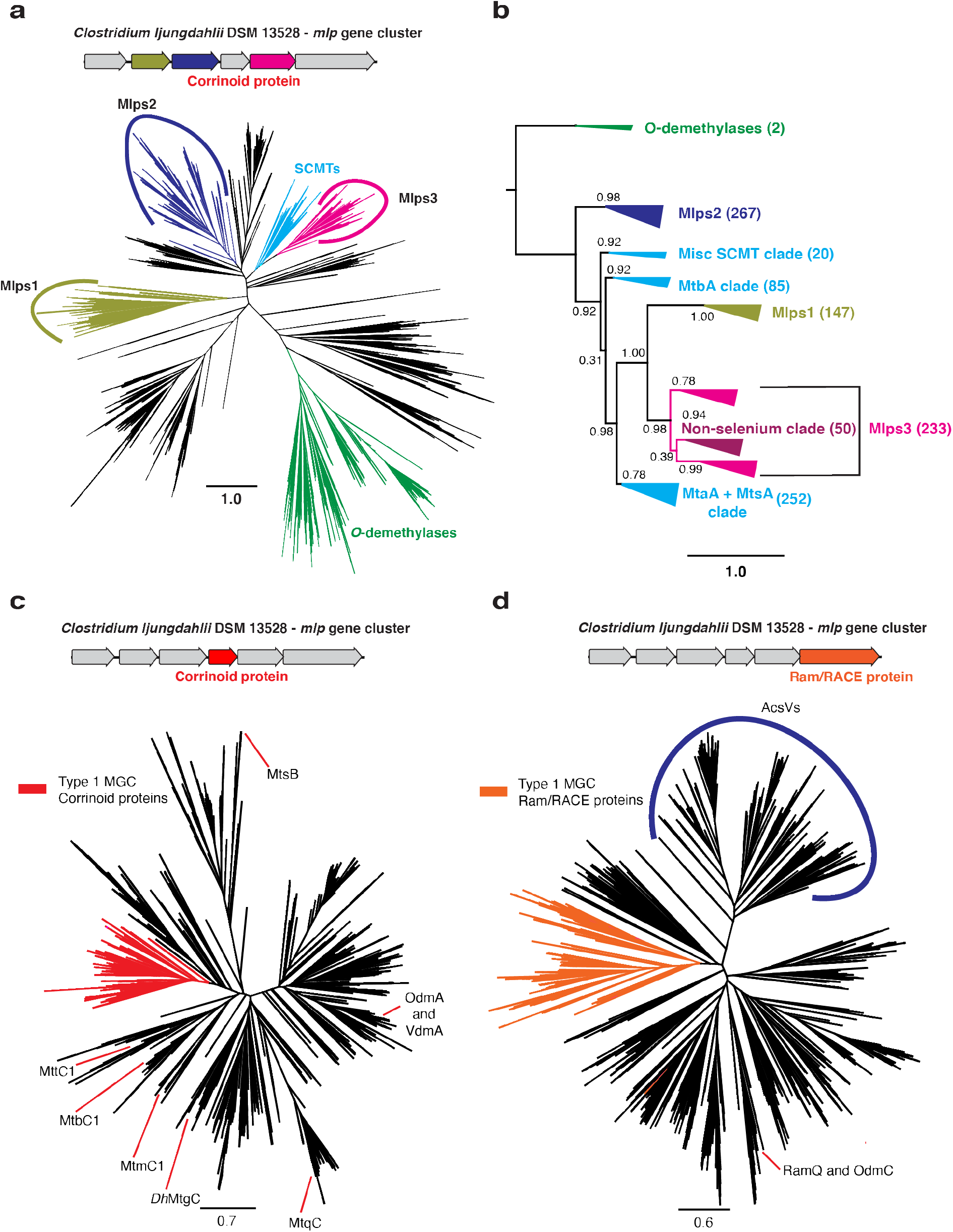
Type I MGCs encode orthologous proteins. **a** An approximately-maximum likelihood (FastTree) phylogenetic tree of members of the UroD superfamily methyltransferases. The FastTree support values for the Mlps1, 2, and 3 clades are 0.95, 0.94, and 1.00, respectively. Scale bar: branch length corresponding to an average of one change per position. **b** A rooted approximately-maximum likelihood (FastTree) phylogenetic tree of selected protein sequences. Shown are the soluble coenzyme M methyltransferases (SCMTs), the Mlps1, 2 and 3, and the phenolic *O-*demethylases OdmB and VdmB (outgroup). Each branch is labeled with the branch’s FastTree support value. The number of sequences in each clade are as follows. *O*-demethylases: 2; Mlps2: 267; Misc. SCMT clade: 20; MtbA clade: 85; Mlps1: 147; Mlps3: 233; MtaA + MtsA clade: 252. **c** An approximately-maximum likelihood (FastTree) phylogenetic tree of members of the methyltransferase corrinoid protein family. The type I MGC proteins are shown in red. The FastTree support value for the type I MGC proteins is 0.92. Selected characterized family members are shown: MtbC1, MtmC1, and MttC1 (all from *M. acetivorans* C2A); *Dh*MtgC from *Desulfitobacterium hafniense* Y51; MtqC from *Eubacterium limosum* ATCC 8486; MtsB from *M. barkeri* Fusaro; OdmA and VdmA from *Acetobacterium dehalogenans* DSM 11527. **d** An approximately-maximum likelihood (FastTree) phylogenetic tree of members of the Ram/RACE protein family. The type I MGC proteins are shown in orange. The FastTree support value for the type I MGC proteins is 0.98. A clade containing the corrinoid iron-sulfur protein reductive activases is labeled “AcsVs”. The locations of the activases RamQ (*E. limosum* ATCC 8486) and OdmC (*A. dehalogenans* DSM 11527) are shown. Scale bars for all tree represent respective branch length corresponding to an average of one change per position.

To further test this hypothesis, we generated two additional phylogenetic trees: one containing the type 1 MGC corrinoid proteins and other monomeric methyltransferase corrinoid proteins (Fig. 3c and 3d, respectively), and the other containing the type 1 MGC Ram/RACE proteins and other Ram/RACE proteins (Fig. S4). In each case, the type 1 MGC proteins were found in the same branch, with a handful of exceptions. These exceptions could be the result of gene conversion or the replacement of a “canonical” type 1 MGC gene with an isofunctional gene from outside of the MGC.

Together, data from the SSN and phylogenetic analyses suggests that type 1 MGCs are homologous in that they encode mutually orthologous proteins. The simplest explanation for the homology of the type 1 MGCs is that they are horizontally transferred as a unit. However, we cannot rule out more complicated scenarios including the transfer of type 1 MGC genes individually into different regions of the genome followed by selective pressure for colocalization. This latter possibility gains some credence when we note the lack of conserved synteny in type 1 MGCs.

We next analyzed the phylogenetic distribution of type 1 MGCs (Fig. 4). To date, we have identified over one hundred prokaryotic genera containing at least one type 1 MGC-encoding strain. Of these, only three (*Methanococcus*, *Methanothermococcus*, and *Methanotorris*) are methanogenic archaea. This is noteworthy because members of the Methanococcales are not known to be methylotrophic and therefore would not be expected to encode SCMTs. All type I MGC-encoding genera only obligate anaerobes, with the majority (73%) belonging to the Firmicutes phylum. Although type 1 MGCs are found a limited number of phyla, they are widely distributed across at least 35 orders of bacteria and archaea. From this analysis, we conclude that type 1 MGCs are phylogenetically widespread and typically associated with microorganisms from anoxic environments. Notable species encoding type 1 MGCs include toxigenic clostridia (*Clostridium botulinum*, *Paeniclostridium sordelii*, certain strains of *Clostridioides difficile*) and industrially-relevant clostridia (*Hungateiclostridium* (*Clostridium*) *thermocellum*, *Clostridium ljungdahlii*).

**Fig. 4.**
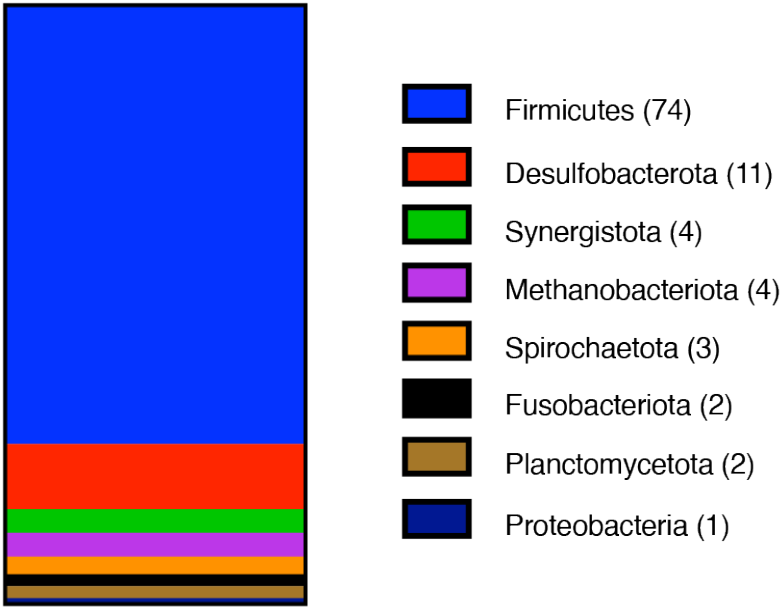
Type I MGCs are found in several prokaryotic Phyla. Phylum composition of genera encoding type I MGCs. 101 genera are represented. The number of type I MGC-encoding genera within each phylum is enclosed in parentheses.

We next analyzed additional components of the type 1 MGCs and proposed a biochemical function for these genes. The UroD superfamily includes corrinoid methyltransferases that (de)methylate heteroatom-containing substrates (10,12,14,15,33). Several lines of evidence suggest that the type 1 MGCs encode a corrinoid *S*- or *Se*-methyltransferase system. First, the closest relationship between known UroD superfamily proteins and the type 1 *mlp* cluster Mlps is with the SCMTs (Fig. a and b), which are *S*-methyltransferases. Additional evidence comes from examination of the enzyme’s Zn^2+^-binding site. This Zn^2+^ cofactor coordinates the heteroatom of a substrate, activating it for (de)methylation (24,33,34). The primary coordination sphere of the Zn^2+^ cofactor is His-Cys-Cys in *S*-methyltransferases from the UroD superfamily, while in *O*-methyltransferases, one or both of the Cys ligands are substituted by carboxylates (Glu or Asp) or are substituted for non-coordinating residues (15,24,33,34). The His-Cys-Cys coordination environment favors the binding of a soft ligand such as a selenium or sulfur atom to the Zn^2+^ center, while substitution of Cys for a carboxylate favors binding of an oxygen atom. The coordination sphere of the type 1 MGC Mlps is (with the exception of several Mlps1) invariably His-Cys-Cys, suggesting that these proteins are *S*- or *Se*-methyltransferases. Homology models of the *C. ljungdahlii* Mlps in further confirmed that these residues are likely appropriately positioned to coordinate Zn^2+^ (Fig. S3). Finally, examination of the non-core components of type I MGCs (genes in type I MGCs that do not encode Mlps, corrinoid proteins, or Ram/RACE proteins) consistently reveals genes involved in sulfur metabolism, including methionine aminotransferases, sulfurtransferases, methionine transporters, and close homologs of the recently-described MarHDK enzyme that catalyzes C–S bond cleavage to produce ethylene from methylthioethanol (35). These observations suggest a potential role for many type I MGCs in methyl-sulfur metabolism.

However, additional evidence suggests that most type I MGCs function in methyl-selenium metabolism. Many type 1 MGCs contain genes provisionally linked to selenium metabolism, such as the recently characterized “dicubane cluster protein” (36–38). Several type 1 MGCs (Fig. 5) contain genes encoding authentic machinery for selenocysteine synthesis, encoding, and decoding (SelABCD), as well as the enzymes of the glycine reductase system, two of which have selenocysteine residues (39,40). Notably, 87% of sequenced species that encode type I MGCs also encode selenocysteine biosynthesis machinery within the same genome. For comparison, only 21.5% of bacteria contain the requisite genes for selenocysteine biosynthesis, and even in the selenocysteine-rich Clostridia, only 42% of 184 genomes encoded selenocysteine biosynthesis machinery (36). Not all organisms that make use of selenium encode selenocysteine biosynthetic machinery—some organisms that use the selenium cofactor only encode selenide-water dikinase (SelD) (36). We therefore manually inspected genomes that contain a type I MGC, but lack SelAB or C, finding that several genomes lacked *selABC*, but had *selD* as well as selenium cofactor-dependent enzymes. As noted below, type 1 MGCs from *C. ljungdahlii* and *C. drakei* are upregulated during autotrophic growth on H_2_ + CO_2_, a condition where there is increased demand for selenoprotein biosynthesis (41). Most importantly, during the preparation of this manuscript, we became aware of prior study that used proteomic and genetic methods to implicate three type I MGC-encoded proteins in the utilization of dimethyl-selenide as a selenium source in the hydrogenotrophic methanogen *Methanococcus voltae* (42). The authors suggested that the Mlps might demethylate (di)methyl-selenide and transfer the methyl group onto the MGC corrinoid protein, liberating selenide for incorporation into biological macromolecules. While the authors did not biochemically characterize the Mlps, their data strongly support a role for the type I MGCs in methyl-selenol utilization.

**Fig. 5.**
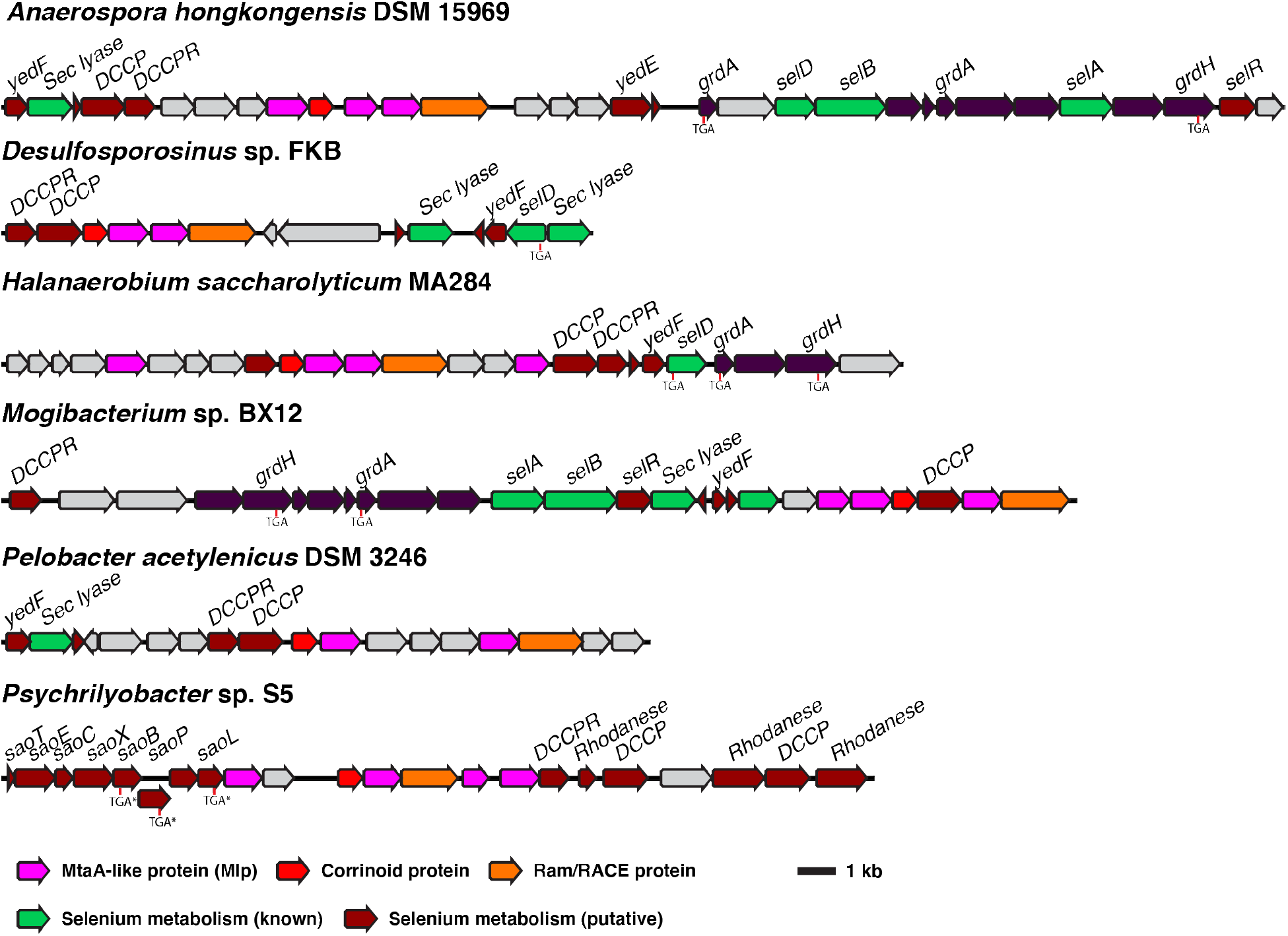
Type I MGCs often encode genes involved in selenium metabolism. Selenium metabolism gene show are either close homologs of genes with experimentally-confirmed roles in selenium metabolism (“Selenium metabolism (known)”) or have been associated with selenium metabolism by bioinformatic methods (“Selenium metabolism (putative)”) (36,37). The positions of selenocysteine codons are make with “TGA.” “TGA*” indicates a selenocysteine codon that has been bioinformatically predicted, but not experimentally confirmed (37). Sec lyase: selenocysteine lyase. DCCP: dicubane cluster protein. DCCPR: dicubane cluster protein reductase. grdA: glycine reductase subunit A. grdH: glycine betaine reductase subunit H. selA: *L*-seryl-tRNA(Sec) selenium transferase. selB: selenocysteine-specific translation elongation factor. selD: selenide water dikinase. selR: putative selenium transcriptional regulator (36). Sao: putative “selenocysteine-assisted organometallic operon genes.

Because type 1 MGCs are present in several model organisms, we examined the expression of these genes in existing proteomic and transcriptomic datasets for *C. botulinum*, *C. ljungdahlii*, and *H. thermocellum* to gain additional information regarding their functions. In all three organisms, the MGCs are expressed at a detectable level under all conditions examined. First, we found that the *C. botulinum* MGC is upregulated during growth at the organism’s optimal growth temperature of 37 °C compared to growth at 15 °C (transcripts 5.27-68.6 fold upregulated at 37 °C) (43). In *C. ljungdahlii*, we found that the type 1 MGC is transcriptionally upregulated roughly 3-5 fold during autotrophic growth on H_2_ + CO_2_ or CO versus heterotrophic growth on fructose (44–46). The same pattern is found in *C. drakei* (6 to 13 fold transcriptional upregulation on H_2_ + CO_2_ compared to fructose) (41). In *H. thermocellum*, the type 1 MGC is upregulated during growth with supplemental acetate in a strain from which the FeFe hydrogenase maturation enzyme *ΔhydG* had been deleted (47). These findings imply that type 1 MGCs are expressed, indicating their physiological relevance, and respond to environmental and genetic perturbations in industrially and medically significant organisms.

Finally, we sought characterize the activity of an Mlp to obtain further insight into the function of type I MGCs. Niess et al.(42) observed that deletion of the gene encoding an Mlp2 (*sdmC*) resulted in loss of the ability of *M. voltae* to use dimethyl-selenide as a selenium source, suggesting the function of Mlps2 may be to demethylate dimethyl-selenide and/or methyl-selenol. We therefore selected *Cl*Mlp2 for biochemical assays. Recombinant ClMlp2 was assayed with sodium selenide and methylcob(III)lalamin under a strictly anoxic nitrogen atmosphere as described in the *Materials and Methods*. In brief, this UV-visible spectroscopy assay assesses methylation of a methyl acceptor by measuring the disappearance or shift of a large methylcob(III)alamin absorbance band (λ_max_ ∼ 525 nm) when the oxidation and/or ligation state of the cobalt ion is changed. For convenience, we assayed for methylcob(III)alamin:selenide methyl transfer (Fig 6a), the reverse of the hypothesized physiological reaction. Addition of sodium selenide (666 μM) to the reaction mixture resulted in a slow but significant decrease in A_525nm_ (Fig. 6b). Control reactions lacking either *Cl*Mlp2 or sodium selenide exhibited rates of A_525nm_ decrease that were 13.7- and 8.3-fold slower, respectively, and are attributable to artefactual photolysis of methylcob(III)alamin inherent to measurement of absorbance (48). Addition of sodium sulfide (666 μM) instead of sodium selenide resulted in a 4.3-fold slower rate, which could suggest the enzyme also possesses low sulfide methylating activity. Starting and ending spectra of the reactions only showed significant spectral changes in the selenide and sulfide reactions. In both cases, the spectra showed a shift in λ_max_ from 520 to 473 nm indicative of formation of base-on cob(II)alamin and therefore demethylation (Fig. 6c). The net steady state formation of cob(II)alamin with minimal accumulation of other intermediates (indicated by a steady absorbance trace at the methylcob(III)alamin/cob(II)alamin isosbestic point of 485 nm (Fig S4)) is likely the result of rapid oxidation of cob(I)alamin, the expected reaction product. Steady-state kinetic analysis revealed an apparent k_cat_ of 0.778 min^-1^ and an apparent K_M_ for selenide of 79.8 μM with a selenide substate inhibition constant (K_I_) of 3830 μM (Fig. 6d). The low k_cat_ is likely due the use of methylcobalamin (rather than the physiological methyl-corrinoid protein) as a methyl donor. The low selenide K_M_, which is comparable to the 46 μM selenide K_M_ measured for *E. coli* selenophosphate synthase (49), suggests that methyl-selenol and/or dimethylselenide are physiological substrates of *Cl*Mlp2. Together, these biochemical studies of *Cl*Mlp2 support the hypothesis that type I MGCs are involved in methyl-selenide metabolism.

**Fig. 6.**
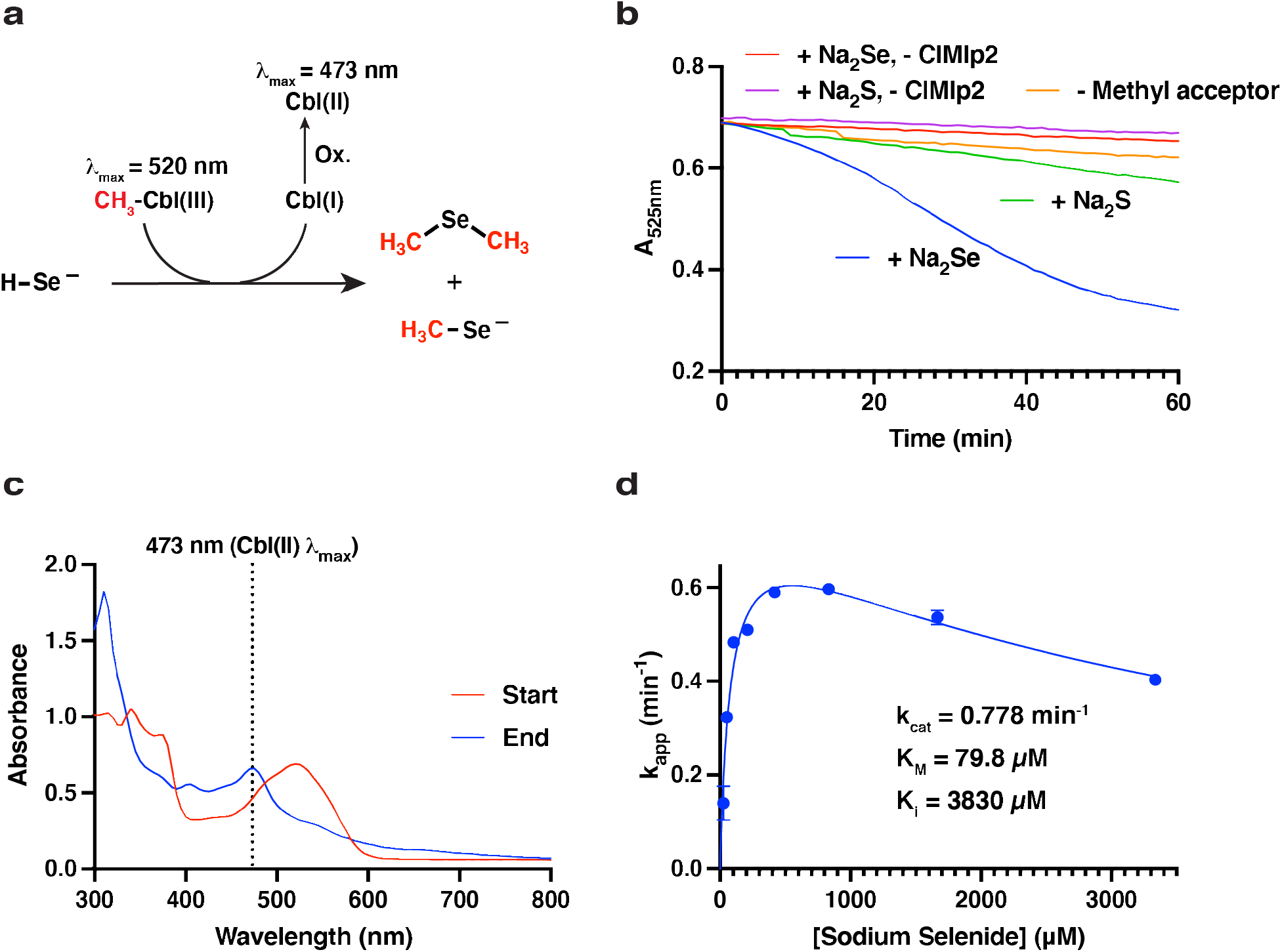
*C. ljungdahlii* (Cl)Mlp2 catalyzes methylcobalamin:selenide methyl transfer. **a** Schematic of the reaction. Cbl: cobalamin. “Ox.” signifies the observed formation of cob(II)alamin, presumably from nonenzymatic reaction of small amounts of diselenides with highly labile cob(I)alamin. **b** 525 nm (methylcob(III)alamin peak) traces indicating disappearance of methylcob(III)alamin. **c** Starting (t = 0 min) and ending (t = 60 min) spectra for the “+ Na_2_Se” reaction shown in **b**. **d** Steady state kinetic analysis of ClMlp2 showing apparent kinetic parameters. Points and error bars represent the mean and standard deviations from three independent reactions.

## Discussion

Methylotrophic methanogenesis is restricted to a subset of the archaea, so it is surprising to find homologs of enzymes from this pathway in non-methanogens. Here, we identify a diversified, widespread set of gene clusters—type 1 MGCs—that encode homologs of the SCMTs involved in methylotrophic methanogenesis. Type I MGCs are found exclusively in anaerobes and are most common in the Firmicutes phylum. There is a strong tendency for type I MGCs to be found in genomes that also contain selenocysteine biosynthesis and decoding machinery.

What reactions do the enzymes encoded by type 1 MGCs catalyze and what is their physiological function? The presence of a corrinoid protein, a corrinoid reductive activase, and homologs of corrinoid methyltransferases (Mlps) implies that the type 1 MGCs encode components of a corrinoid methyltransferase pathway. Protein sequence comparisons indicate that the Mlps are likely *S*- or *Se*-methyltransferases that (de)methylate thiol and selenol metabolites.

The potential physiological roles of such reactivity may be explored through considering features of the organisms that possess type I MGCs. Earlier studies clearly implicate type I MGCs in methyl-selenol valorization (39). However, those authors examined only the corrinoid protein and two closely-related Mlps2 in dimethyl-selenide utilization—the more distantly related *M. voltae* Mlp1 and Mlp3 appear to have gone unnoticed. It seems likely that the full type I MGC contains methyltransferases for demethylation of several methyl-selenium compounds. While the genetic data in Ref. 39 implicates dimethyl-selenide and/or methyl-selenol as substates of the type I MGC Mlps2 (an inference that our assay of *Cl*Mlp2 supports), we propose that the Mlps1 and Mlps3 may be responsible for demethylation of other methyl-selenium compounds such as methyl-selenocysteine and selenomethionine and concomitant methylation of the gene cluster’s encoded corrinoid proteins. We propose that the methyl-selenide-derived methyl group is subsequently transferred from the methylated corrinoid protein to a thiol-containing compound such as sulfide. The resulting methyl-sulfur metabolite could then be excreted or recycled into methionine. Forms of methyl-selenium are present in many environments, including the human gut (50). It may be that most type I MGCs function to valorize these methyl-selenium compounds. However, many organisms (such as *C. ljungdahlii* and *M. voltae*) upregulate their type I MGCs even when methyl-selenium compounds are not added to growth media. This observation suggests that there may be an endogenous source of methyl-selenium compounds. Indeed, it has been suggested and evidentially supported that the selenide generated as an intermediate in selenoprotein biosynthesis can react non-enzymatically with *S*-adenosylmethionine (SAM) (51). This presents a problem for a cell that has a high demand for selenium metabolism: methyl selenide formation from SAM wastes valuable selenium and generates a potentially toxic dead-end metabolite. The enzymes encoded by type I MGCs may solve this quandary by recycling the methyl selenide into methionine.

Some organisms, such as *H. thermocellum*, encode and express type I MGCs but lack genes encoding known selenoenzymes or other indicators of selenium metabolism. We found that all the selenium-trait deficient organisms have Mlps3 that fall into a single clade of the Mlp phylogenetic tree (Fig. 3b). We suggest that the corresponding type I MGCs are involved in methyl-sulfur metabolism. While one group of type I MGCs (>90%) is likely involved in methyl-selenium metabolism, the other, which encodes Mlps3 from the non-selenium clade, probably has a selenium-independent function.

Evidence from genomic context suggests that the non-selenium MGCs could be involved in methionine salvage, defined as a metabolic pathway that converts methylthio-adenosine (MTA) into methionine (52). MTA is a toxic by-product of the biosynthesis of key metabolites derived from SAM (53). Methionine salvage pathways therefore support sulfur metabolism both by detoxifying a by-product and preserving the metabolically expensive methylthiol group of MTA (54). The so-called “universal” methionine salvage pathway is O_2_-dependent, and methionine salvage pathways that operate under anaerobic conditions have only recently been discovered (52). The anaerobic methionine salvage pathway from *Rhodospirillum rubrum* was recently shown to use the methylthio-alkane reductase MarHDK system, homologs of the nitrogenase subunits NifHDK, respectively (35). Remarkably, *MarHDK*-like genes belonging to the nitrogenase superfamily IV-A group are prevalent in or near type I MGCs from the non-selenium clade and, when not present in the type I MGC itself, are invariably present elsewhere in the genome (Fig. 7a). This suggests that the non-selenium type I MGCs encode enzymatic machinery for either methionine salvage, sulfur valorization from methyl-sulfide compounds, or both.

**Fig. 7.**
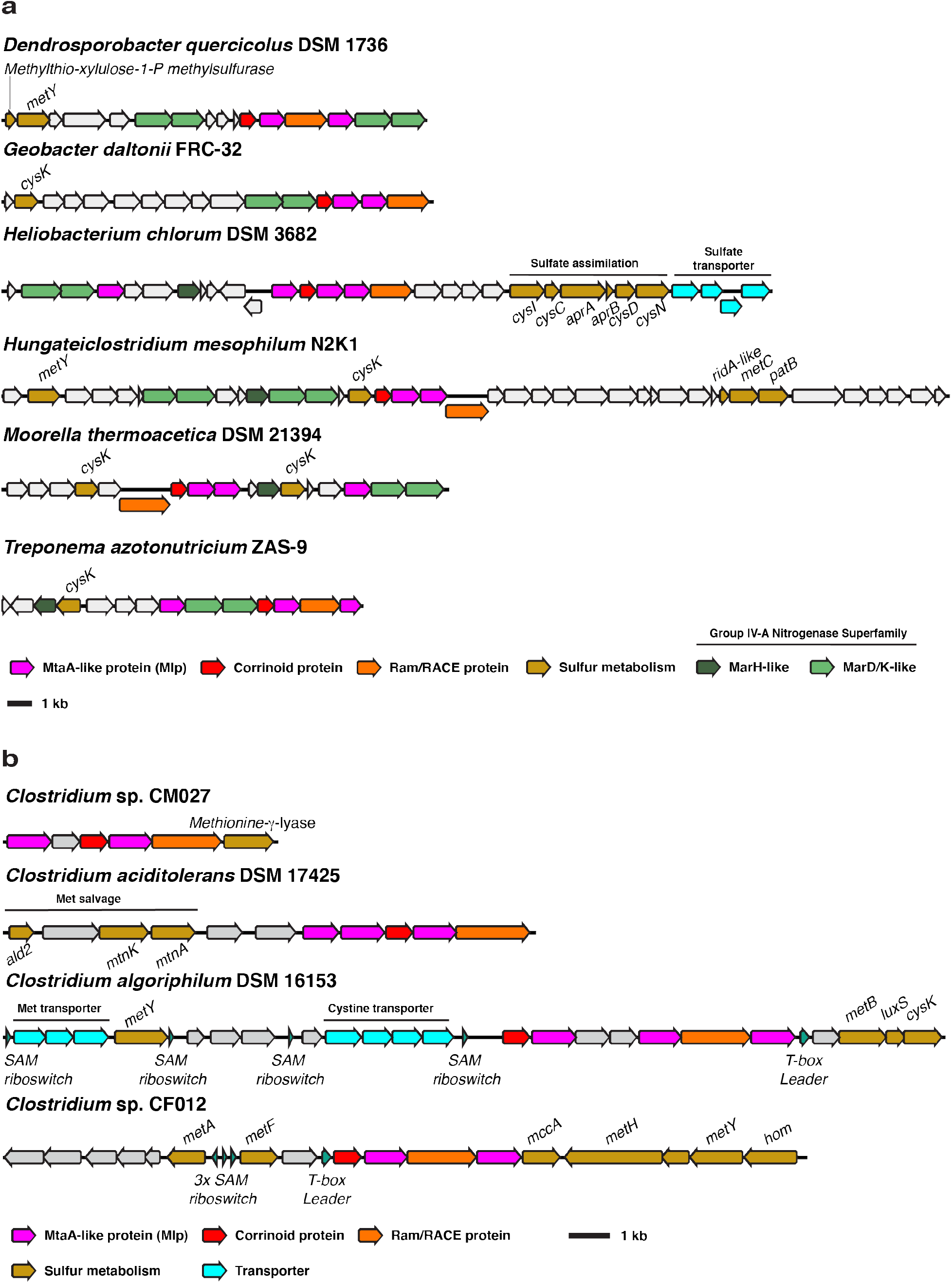
Type I MGCs from encoding an Mlp3 from the non-selenium clade also encode putative sulfur and methionine metabolism genes. **a** Non-selenium type I MGCs associated with group IV-A nitrogenase superfamily genes and other genes involved in sulfur metabolism. Methylthioribulose-1-P methylsulfurase is a methionine salvage enzyme originally described in *Rhodospirillum rubrum* (55). **b** Other non-selenium type I MGCs associated with methionine metabolism. MetY: *O*-acetyl-homoserine sulfhydrylase. CysK: cysteine synthase. CysI: assimilatory sulfite reductase. CysC: adenylyl-sulfate kinase. AprAB: phosphoadeninosine-phospho-sulfate reductase subunits A and B. CysND: sulfate adenylyltransferase subunits 1 and 2. MetC cystathionine beta-lyase. PatB: alternate cystathionine beta-lyase. Ald2: methylthio-ribulose-1-phosphate aldolase. MtnK: methylthio-ribose kinase. MtnA: methylthio-ribose-1-phosphate isomerase. MetB: cystathionine gamma-synthase. LuxS: *S*-ribosyl-homocysteine lyase. MetA: homoserine-*O*-succinyltransferase. MetF: methylene-tetrahydrofolate reductase. MccA: *O*-acetyl-serine dependent cystathionine beta-synthase. MetH: cobalamin-dependent methionine synthase. Hom: homoserine dehydrogenase. SAM: *S*-adenosyl-methionine.

Additionally, the sporadic distribution of type I MGCs across phylogenetic trees is potentially consistent with a role in methionine salvage, as such pathways are dispensable under many growth conditions (54) and are not strictly conserved even in closely related strains (56).

We propose how a type I MGC-encoded methionine salvage pathway could operate in Fig. 8, considering a type I MGC that encodes only two Mlps. The MarHDK-like enzymes may have a role in such a pathway, generating methyl-sulfide that might serve as the methyl donor for the MGC methyltransferase system (R^1^-SCH_3_ with R^1^ = H). Alternatively, the MarHDK-like proteins might serve as part of a second, parallel methionine salvage pathway, or even as a branch of a topologically bifurcating pathway with two routes to methionine.

**Fig. 8.**
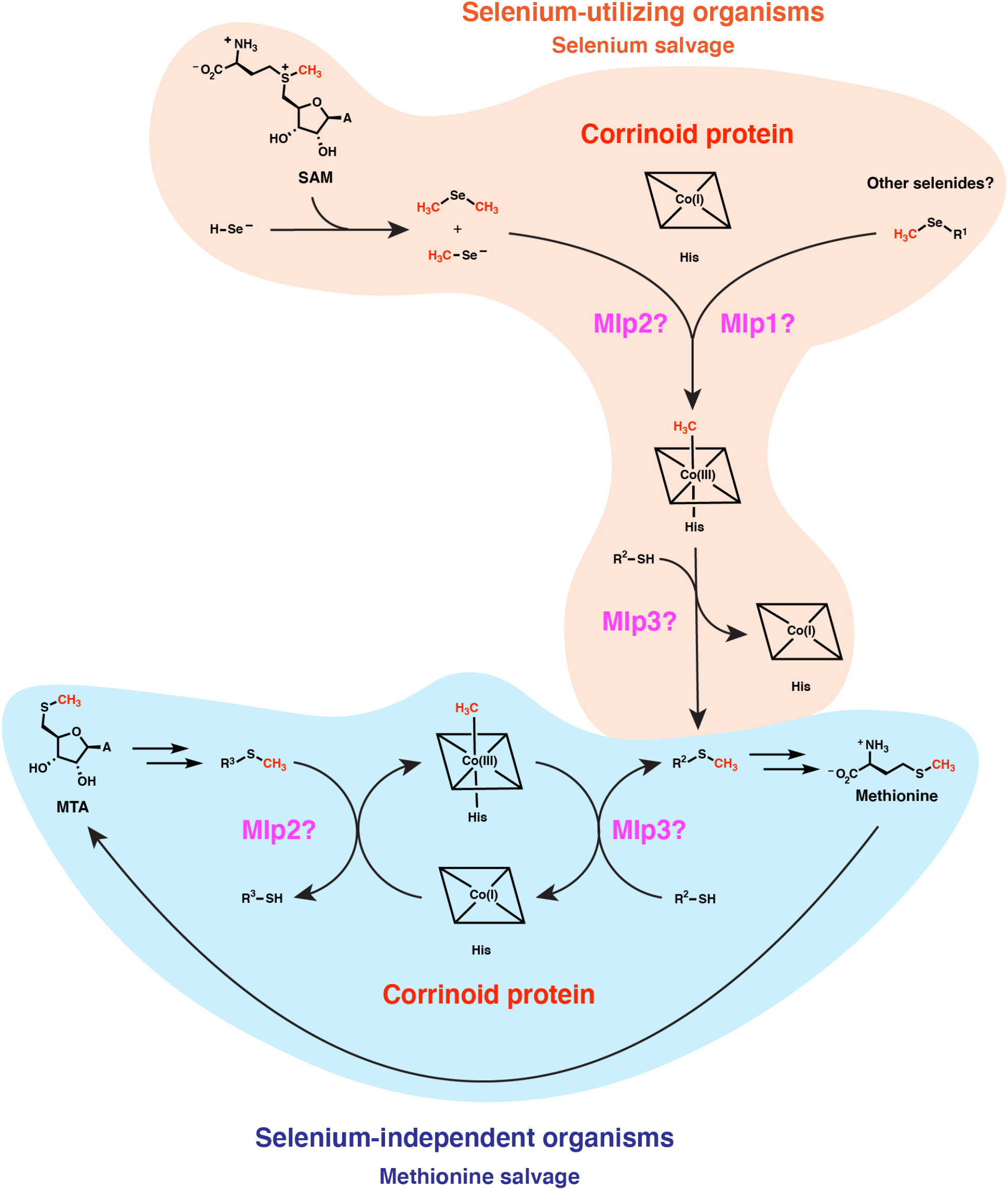
Type I MGCs likely encode components of corrinoid-dependent selenium salvage or methionine salvage pathways. The vast majority of type I MGC encoders likely use these methyltransferase systems to recycle methylated selenides formed from the non-enzymatic reaction of SAM with selenol compounds, potentially recycling the methyl group into methionine. A smaller subset of type I MGC encoders in the ref seq database (about 10%) lack known selenium metabolism traits. We propose that these MGCs have evolved to participate in a methionine salvage pathway. MTA: 5’-methylthio-adenosine. His: histidine. A: adenine.

Intriguingly, this proposed methionine salvage pathway uses a distinct biochemical logic from related pathways. Characterized methionine salvage pathways all incorporate the methylthiol group of MTA intact into methionine (35,52,53,56). In the proposed methionine salvage pathway involving the type 1 MGCs, the methylthio-ether of MTA is dismantled into a methyl group which is carried by the corrinoid protein, leaving the free thiol behind. This pathway architecture theoretically avoids production of volatile methyl-sulfur intermediates that could be more readily lost from the cell. The proposed role of a corrinoid cofactor in methionine salvage is also novel. Ultimately, biochemical characterization of the enzymes encoded by type I MGCs will be required to test this proposal.

In summary, we have identified a class of microbial gene clusters encoding components of probable corrinoid *S*- and *Se*-methyltransferase pathways. The type I MGCs are found in phylogenetically and metabolically diverse organisms encompassing >100 microbial genera. Bioinformatic analyses and preliminary biochemical studies suggest type I MGCs participate in methyl-selenide metabolism and/or methionine salvage. Ultimately, this previously unappreciated example of methanogen specific enzymes encoded outside of the archaea could provide insights into the vitally important fields of sulfur and selenium cycling.

## Supporting information

Supplemental Tables and figuresental

Phylogenetic tree data

List of species with Type I MGCs

## Acknowledgements

D.J.K. acknowledges support from National Institutes of Health Training Grant #5T32GM095450. This work was supported by a Howard Hughes Medical Institute (HHMI)-Gates Faculty Scholar Award (OPP1158186) and a National Science foundation (NSF) Alan T. Waterman Award (CHE-20380529) to E.P.B. E.P.B. is Howard Hughes Medical Institute Investigator. We also note that this article is subject to HHMI’s Open Access to Publications policy. HHMI lab heads have previously granted a nonexclusive CC BY 4.0 license to the public and a sublicensable license to HHMI in their research articles. Pursuant to those licenses, the author-accepted manuscript of this article can be made freely available under a CC BY 4.0 license immediately upon publication.

## Data Availability Statement

All relevant data has been made available in the main text, Supplementary Information, and associated data files.

