## Supplemental Tables and figuresental for "A diversified, widespread microbial gene cluster encodes homologs of methyltransferases involved in methanogenesis"

### for

#### Supplementary methods

Construction of homology models

Clostridium ljungdahlii DSM 13528 Mlps 1, 2, and 3 were submitted to the SWISS-MODEL webserver (23) using *Methanosarcina mazei* MtaA (PDB ID: 4AY7) (24).

Expression and purification of *Clostridium ljungdahlii* MtaA-like protein 2 (*Cl*Mlp2)

*Clmlp2* gene was PCR-amplified from *C. ljungdahlii* DSM 13528 genomic DNA and cloned using the PIPE method into pSpeedET, which was then transformed into *E. coli* Rosetta 2(DE3)pLysS. The resulting strain was grown aerobically at 37 °C with 180 RPM shaking in four 4 L baffled flasks, each containing 2 L of LB broth (Research Products International) supplemented with 22 mM potassium phosphate (salts from Sigma Aldrich) buffer pH 7.2, 20 mM D-glucose (Sigma Aldrich), 20 mM $MgCl_2$ (Sigma-Aldrich), 50 μg/mL kanamycin sulfate (VWR International), and 35 μg/mL chloramphenicol (Sigma Aldrich). At a $OD_{600nm}$ of 0.6, IPTG (Teknova) was added to a final concentration of 0.5 mM, and the growth temperature was shifted to 16 °C. After 12 hours of incubation at 16 °C, the cultures were harvested by centrifugation and the resulting pellets washed once with 50 mM MOPS (Sigma Aldrich) pH 7.2 before resuspension in buffer A. Buffer A contained

50 mM imidazole, 22 mM potassium phosphate, 500 mM NaCl, and 10% glycerol (by volume). The resuspended pellet was lysed by sonication using a Branson Digital Sonifier in 120 cycles composed of 5 seconds of sonication followed by 15 seconds of rest. The resulting lysate clarified by centrifugation at 50,000 x g for 45 minutes. The extract was loaded onto a 5 mL Hitrap IMAC FF $Ni^{2+}$-affinity column (GE Life Sciences), which was washed with buffer A. The elution was conducted with a linear gradient from 100% buffer A to 100% buffer B. Buffer B contained 500 mM imidazole, 22 mM potassium phosphate, 500 mM NaCl, and 10% glycerol. The eluted protein was collected, concentrated with a 5 kDa-cutoff Spin-X UF concentrator (Corning), and diluted at least 10-fold in buffer C. Buffer C consisted of 50 mM MOPS, 10% glycerol, pH 7.2. The resulting protein sample was loaded onto a 5 ml Hitrap Q column, which was washed with buffer C, followed with elution with a linear gradient from 100% buffer C to 100% buffer D. Buffer consisted of 50 mM MOPS, 500 mM NaCl, 10% glycerol, pH 7.2. The resulting fractions were screened for a polypeptide of the expected molecular mass (39.1 kDa) for *Cl*Mlp2. The identified fractions were concentrated and buffer exchanged into buffer E (50 mM MOPS, 200 mM KCl, 10% glycerol, pH 7.2). The resulting protein preparation was made anoxic under an $N_2$ atmosphere by the flush-evacuation technique using a Schlenk line. It was then flash frozen at in liquid $N_2$ and stored at –70 °C until use.

Assay of *Cl*Mlp2

The biochemical assay procedure used was described previously (25). The assays were conducted in a Powerwave XS (BioTek) plate reader heated to 37 °C in a quartz 96-well plate (Molecular Devices). All phases of the assay took place under red light and otherwise in the dark in an MBraun chamber under a 100% $N_2$ atmosphere. 150 μL reaction mixtures (assembled in the plate and without methyl acceptor) were incubated at 37 °C for 2 minutes and were then initiated with addition of methyl acceptor substrate. Following addition of the methyl acceptor, the reaction composition was 50 mM MOPS pH 7.2, 0.4 mM methylcobalamin (Sigma Aldrich), 4.7 μM *Cl*Mlp2, and methyl acceptor at indicated concentrations. Following initiation, the absorbances at 525, 485, and 800 nm were followed. Rates of reaction were calculated from the maximum rate of $A_{525nm}$ decrease sustained over at least 5 minutes during the 1 hour period for which the reaction was followed. The absorbance change rate was then converted using the $\Delta\varepsilon_{525nm}$ = 8.6 $mM^{-1}$ reported in (25). Manipulations and analyses of the data were performed in GraphPad Prism 9.3.1.

<u>Phylogenetic membership of type I MGC-encoding organisms</u>

We analyzed the phylogenic composition of type I MGC-encoding bacteria and archaea according to the Genomes Taxonomy Database (GTDB) taxonomic system (22).

| Protein name | Annotation | Locus tag | E value | % amino acid identity |
|---|---|---|---|---|
| *Cl*Mlp3 | methylcobamide--CoM methyltransferase | CLJU_RS09375 | 4.58e-60 | 32.5 |
| *Cl*UroD7 | methylcobamide--CoM methyltransferase | CLJU_RS10340 | 1.55e-54 | 32.2 |

| | | | | |
|---|---|---|---|---|
| *Cl*UroD1 | uroporphyrinogen decarboxylase | CLJU_RS10345 | 2.29e-38 | 30.7 |
| *Cl*UroD8 | methylcobamide--CoM methyltransferase | CLJU_RS10365 | 8.59e-32 | 27.6 |
| *Cl*UroD9 | methylcobamide--CoM methyltransferase | CLJU_RS14945 | 7.10e-29 | 27.0 |
| *Cl*Mlp2 | uroporphyrinogen decarboxylase | CLJU_RS09365 | 9.67e-26 | 27.0 |
| *Cl*Mlp1 | methylcobalamin--coenzyme M methyltransferase | CLJU_RS09360 | 2.57e-10 | 25.8 |

**Table S1** The results of BLASTp search against the *C. ljungdahlii* DSM 13528 predicted proteome using *M. barkeri* Fusaro MtbA (uniprot ID: O30640) as a query.

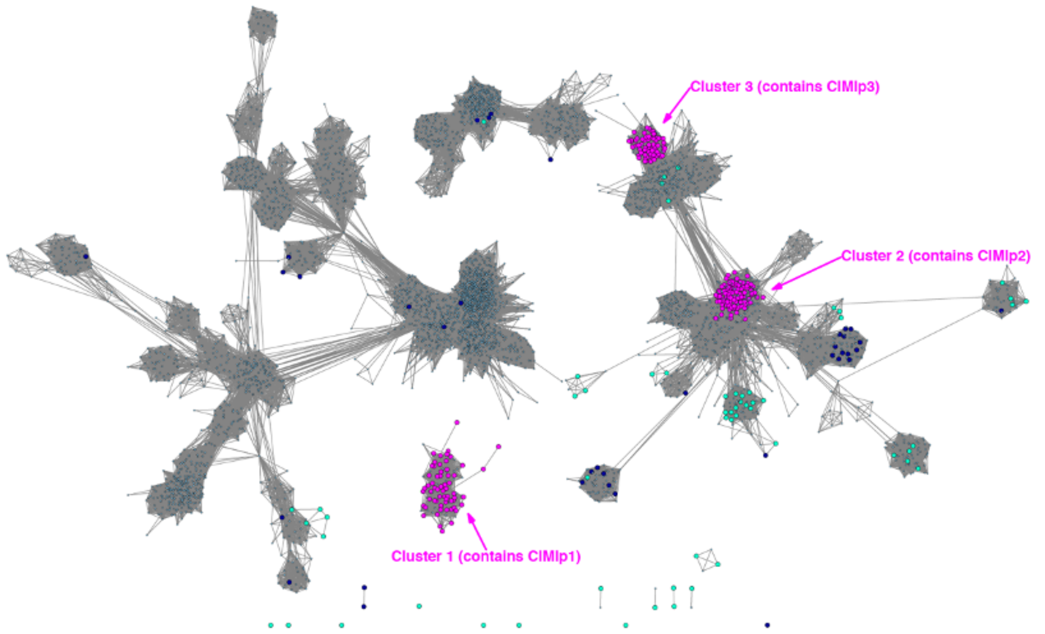


**Fig. S1** Protein SSN of the UroD superfamily (IPR000257). The alignment score cutoff was 40 and the node representation percent cut was 50 % aa identity. Only nodes that are directly or indirectly connected to MGC Mlps are shown for simplicity. Magenta: core type 1 MGC Mlps. Cyan: “auxiliary” type 1 MGC cluster Mlps. Dark Blue: Mlps from non-type 1 MGCs.

**a**

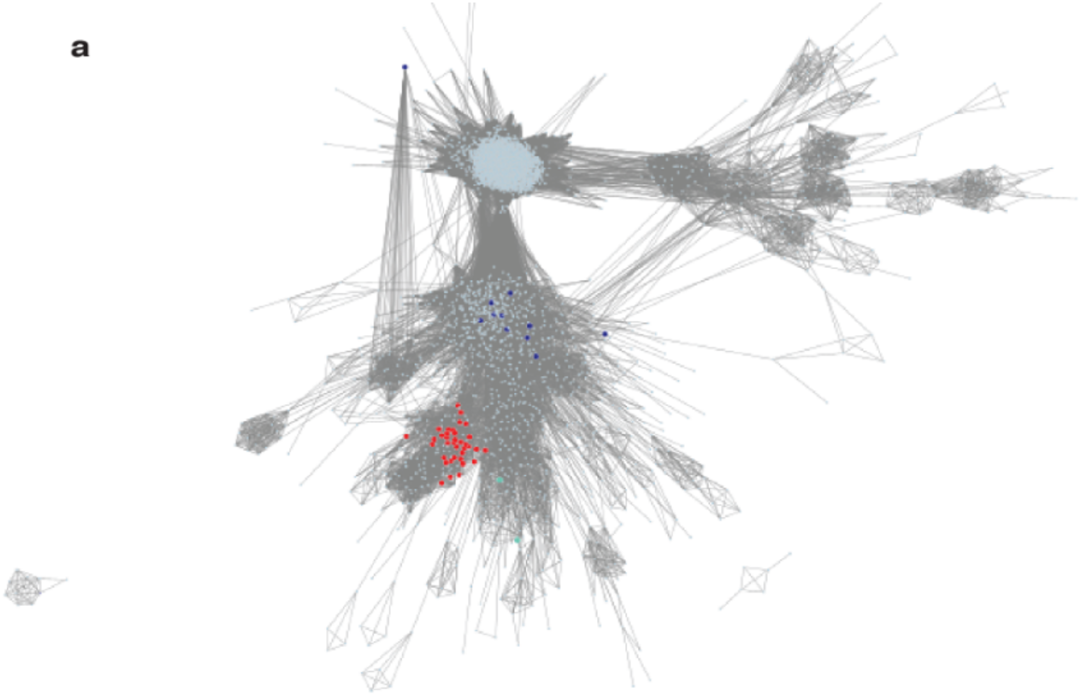

**b**

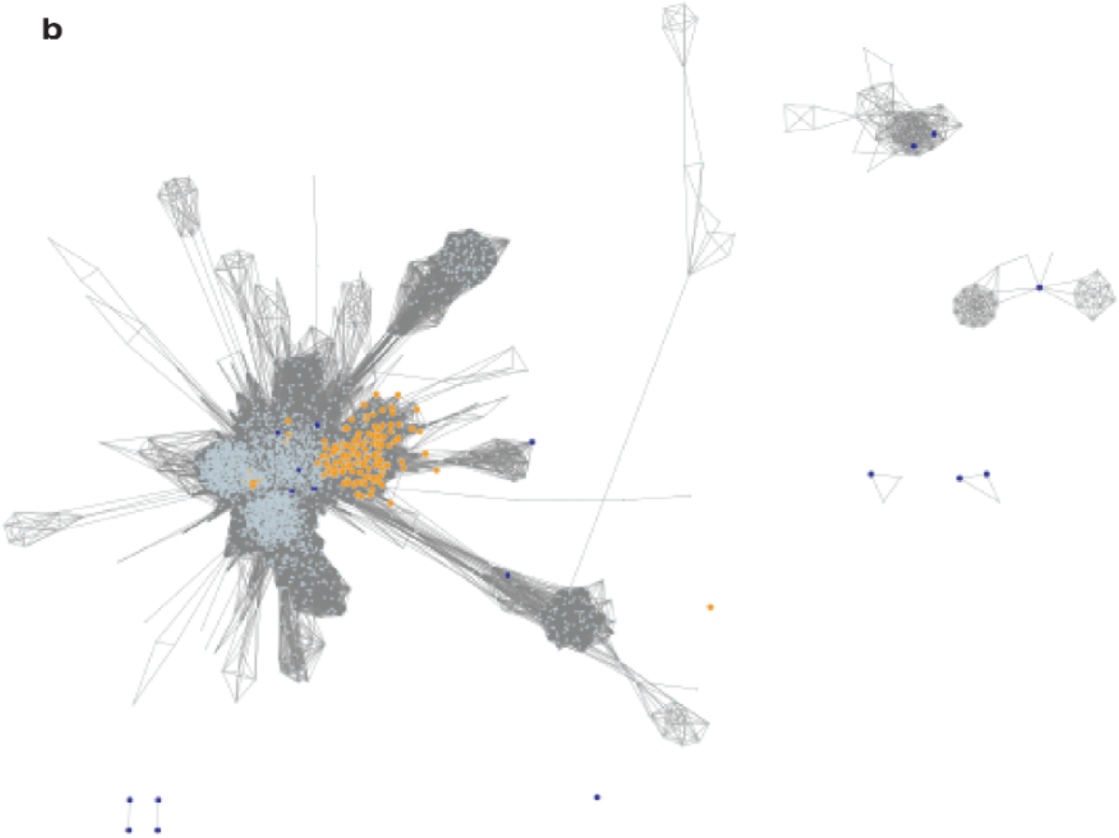

**Fig. S2 a** Protein SSN of the corrinoid-binding superfamily (IPR006158). The alignment score cutoff was 50 and the node representation percent cut was 50 % aa identity. Only nodes that are directly or indirectly connected to MGC corrinoid proteins are shown for simplicity. Magenta: canonical type 1 MGC corrinoid proteins. Cyan: “auxiliary” type 1 MGC cluster corrinoid proteins that are rarely present in a MGC that already has a canonical corrinoid protein. Dark Blue: corrinoid proteins from non-type 1 MGCs. **b** Protein SSN of the “bacterial-type” Ram/RACE superfamily (IPR027980). The alignment score cutoff was 100 and the node representation percent cut was 50 % aa identity. Only nodes that are directly or indirectly connected to MGC Ram/RACE proteins are shown for simplicity. Magenta: canonical type 1 MGC Ram/RACE proteins. Dark Blue: corrinoid proteins from non-type 1 MGCs.

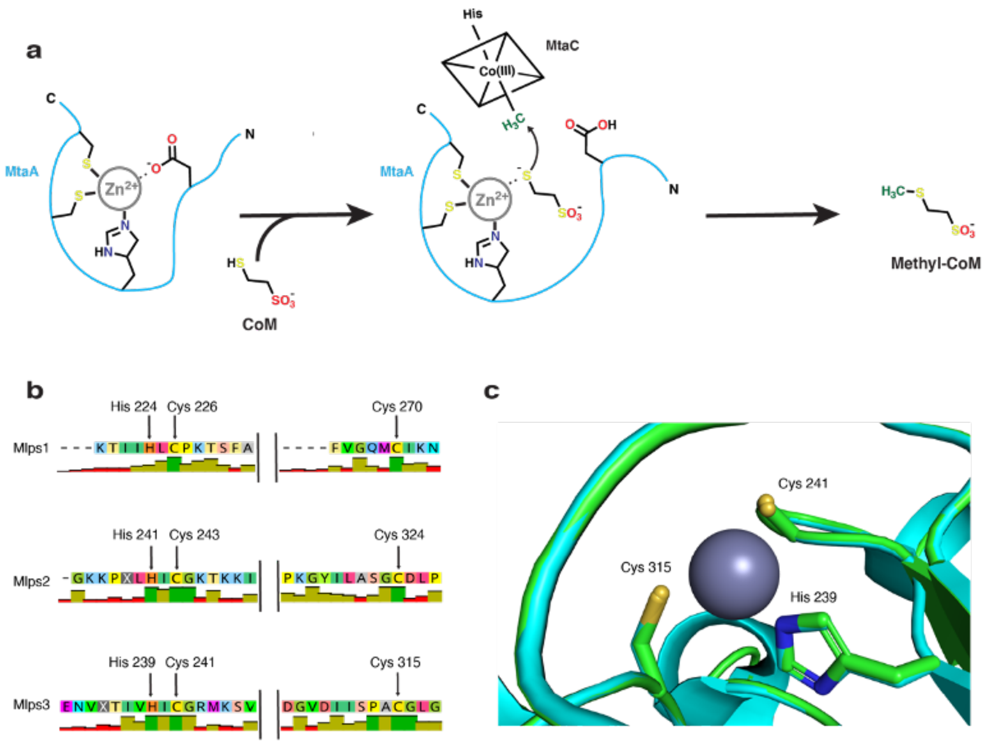


**Fig. S3 Sequence alignments and homology models of the type I MGC Mlps reveal a conserved $Zn^{2+}$ binding site. a** The catalytic role of $Zn^{2+}$ in MtaA and other SCMTs. Coenzyme M (CoM) displaces a conserved carboxylate ligand (Asp or Glu) and is deprotonated, activating the CoM thiolate for nucleophilic attack on the methylated corrinoid protein (MtaC). N and C represent the MtaA N- and C-termini, respectively. **b** Alignments of the Mlps with the positions of the $Zn^{2+}$ ligands indicated with black arrows. The height and color of the bars indicates the extent of amino acid conservation. Tall bars correspond to strongly conserved amino acids, while short bars correspond to low conservation. Alignments were constructed from 147, 268, and 233 sequences encoding Mlps1, 2, and 3, respectively. Green bars represent a completely conserved amino acid while yellow and red bars represent moderately and un-conserved amino acids respectively. Consensus sequences present as simple majority residue representation. **c** A homology model of *Cl*Mlp3 (green) showing the appropriate positioning of the $Zn^{2+}$ (grey sphere) cofactor. *Methanosarcina mazei* MtaA (PDB ID: 4ay7, cyan) was used as a template and is overlayed (cyan). In **b** and **c**, the residue numbering for the *C. ljungdahlii* proteins is shown.

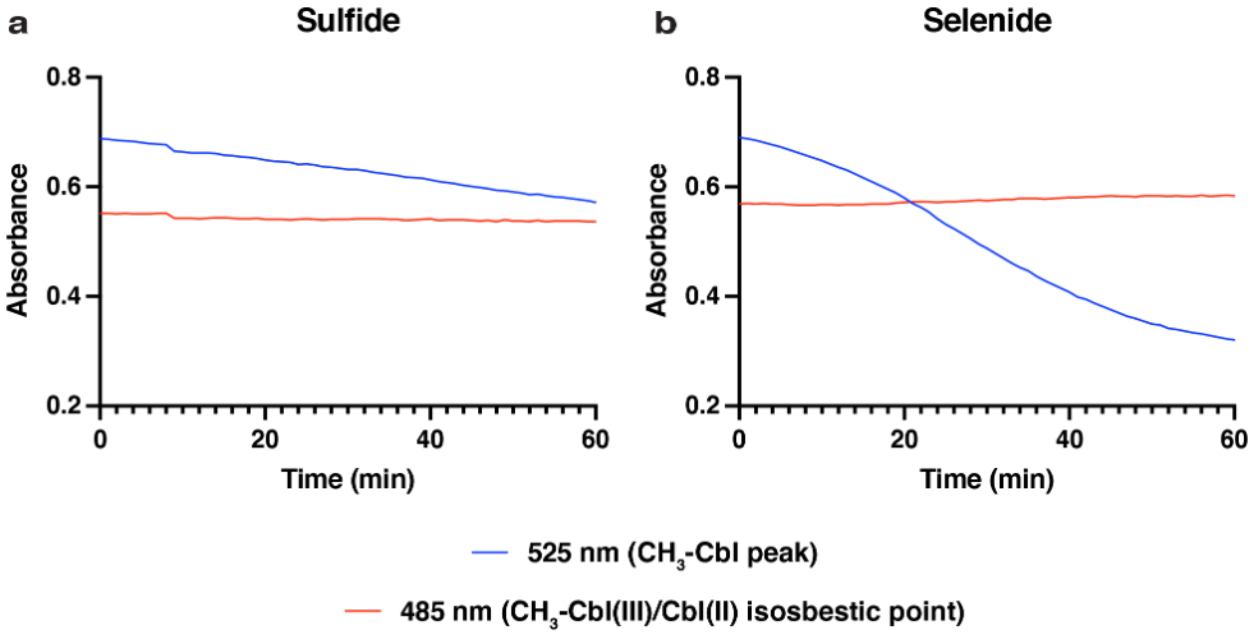


**Fig. S4 Absorbance traces of the *Cl*Mlp2 reactions showing steady-state conversion of methylcob(III)alamin to cob(II)alamin with insignificant accumulation of other intermediates. a** with sodium sulfide as a methyl acceptor. **b** with sodium selenide as a methyl acceptor. $CH_3$-Cbl(III): methylcob(III)alamin. Cbl(II): cob(II)alamin. The reactions are the same as those in Fig 6b.
